# BDNF/TrkB.T1 signaling is a novel mechanism for astrocyte morphological maturation

**DOI:** 10.1101/518787

**Authors:** Leanne M. Holt, Natasha L. Pacheco, Raymundo Hernandez, Muhannah Hossain, Michelle L. Olsen

**Affiliations:** Department of Cell, Developmental, and Integrative Biology, University of Alabama at Birmingham, 1918 University Blvd., Birmingham, Alabama 35294; School of Neuroscience, Virginia Polytechnic and State University, Life Sciences Building Room 213, 970 Washington St. SW, Blacksburg, Virginia 24061.

**Author notes:** Address correspondence to: Michelle Olsen, PhD, School of Neuroscience, 970 Washington Street SW, LS1 RM 213, Virginia Polytechnic and State University, Blacksburg, Virginia, 24060.

**Keywords:** astrocyte, BDNF, TrkB, development, synaptogenesis, morphogenesis

## Abstract

Brain derived neurotrophic factor (BDNF) is a critical growth factor involved in the maturation of neurons, including neuronal morphology and synapse refinement. Herein, we demonstrate astrocytes express high levels of BDNF’s receptor, TrkB (in the top 20 of protein-coding transcripts), with nearly exclusive expression of the truncated isoform, TrkB.T1 which peaks in expression during astrocyte morphological maturation. Using a novel culture paradigm, we show that astrocyte morphological complexity is increased in the presence of BDNF and is dependent upon BDNF/TrkB.T1 signaling. Deletion of TrkB.T1 *in vivo* revealed morphologically immature astrocytes with significantly reduced volume and branching, as well as dysregulated expression of perisynaptic genes associated with mature astrocyte functions, including synaptogenic genes. Indicating a role for functional astrocyte maturation via BDNF/TrkB.T1 signaling, TrkB.T1 KO astrocytes do not support normal excitatory synaptogenesis. These data suggest a significant role for BDNF/TrkB.T1 signaling in astrocyte morphological maturation, a critical process for CNS development.

## Introduction

Astrocyte maturation is crucial developmental processes for normal CNS function. In the rodent cortex, astrocyte maturation takes place largely during the first 2–4 postnatal weeks. Importantly, this includes morphological maturation wherein immature astrocytes elaborate their processes and infiltrate the neuropil with fine, terminal, leaflet processes (Bushong et al., 2004). These leaflet terminals represent important functional structures, allowing cell-cell communication with neighboring astrocytes and enwrapping of synapses—where astrocytes participate in neurotransmitter uptake and synapse development and stabilization (Farhy-Tselnicker and Allen, 2018; Oberheim et al., 2012). Underscoring the morphological complexity of these cells, estimates indicate a single mature rodent astrocyte encompasses between 20,000 – 80,000 μM^3^ of domain space (Bushong et al., 2002; Halassa et al., 2007), associates with 300-600 neuronal dendrites (Halassa et al., 2007), and contacts more than 100,000 individual synapses (Freeman, 2010). The maturation period of astrocyte morphogenesis coincides with neuronal synaptic refinement (Freeman, 2010; Morel et al., 2014) and differential expression of key genes associated with mature astrocyte functions, such as Glt1, Kir4.1, and Aqp4 (Clarke et al., 2018; Molofsky and Deneen, 2015; Molofsky et al., 2012; Morel et al., 2014; Nwaobi et al., 2014). While the time course of astrocyte morphogenesis is well defined, few studies have attempted to identify molecular signals guiding astrocyte morphogenesis and maturation. To date, three mechanisms have been identified: Fibroblast Growth Factor (FGF)/Heartless signaling (Stork et al., 2014), glutamate/mGluR5 signaling (Morel et al., 2014), and contact-mediated neurexin/neuroligin (Stogsdill et al., 2017).

BDNF (Brain Derived Neurotrophic Factor) is a critical growth factor in the development, maturation, and maintenance of the CNS. Its role in neuronal cell growth, differentiation, morphology, and synaptogenesis via TrkB receptor signaling is well characterized (Autry and Monteggia, 2012; Fenner, 2012; Park and Poo, 2013). In the CNS, TrkB has two main isoforms. The full-length receptor, TrkB.FL, possesses a tyrosine kinase domain that autophosphorylates with BDNF binding, and a truncated receptor, TrkB.T1. While TrkB.T1 lacks the canonical tyrosine kinase domain, BDNF binding to this receptor is thought to signal through a RhoGTPase inhibitor and the phospholipase C (PLC) pathway (Deinhardt and Chao, 2014; Fenner, 2012). Dysregulation of BDNF/TrkB signaling has been implicated in multiple neurological and neurodevelopmental disorders (Park and Poo, 2013). However, a role for BDNF in the developmental maturation of astrocytes has not been investigated.

Here for the first time we demonstrate that *Ntrk2*, the gene that encodes BDNF’s receptor is highly enriched in astrocytes, particularly during the critical period of astrocyte morphological maturation. RNA sequencing and qPCR reveal that astrocytes predominately express the truncated TrkB (TrkB.T1) receptor. TrkB.T1 receptor expression mediates increased astrocyte morphological complexity in response to BDNF *in vitro*, and TrkB.T1 KO astrocytes *in vivo* remain morphologically immature with significantly reduced cell volumes and morphological complexity. TrkB.T1 KO astrocytes exhibit dysregulation of genes associates with perisynaptic mature astrocyte function, including synaptogenic genes. Finally, co-culture studies indicate TrkB.T1 KO astrocytes do not support normal synaptic development. Together, these data suggest a significant role for BDNF/TrkB.T1 signaling in astrocyte morphogenesis and indicate this signaling may contribute to astrocyte regulation of neuronal synapse development.

## Results

### Astrocytes express high levels of truncated TrkB.T1 mRNA

RNA sequencing was performed on cortical astrocytes that were acutely isolated from late juvenile (PND 28) mice via magnetic separation (n = 5 replicates) (Holt and Olsen, 2016). Postnatal day 28 was chosen based on reports indicating astrocytes are considered morphologically mature at this stage of development (Bushong et al., 2004; Morel et al., 2014). Analysis of this sequencing data revealed *Ntrk2*, the gene encoding BDNF’s high affinity receptor TrkB, to be in the top 20 of all protein coding RNA’s (#18) detected. The two isoforms are distinguishable given that the full-length receptor contains exons for the tyrosine kinase domain, while the truncated TrkB.T1 receptor lacks this domain but has an additional exon (exon 12) not found in the full length receptor (Fig. 1A). Therefore, isoform-specific transcript expression was analyzed from the isolated astrocytes and corresponding whole cortex. This analysis revealed that PND28 astrocytes predominately express the truncated isoform, with nearly 90% of all *Ntrk2* expression in cortical astrocytes attributed to TrkB.T1 (151.91 +/− 9.18 FPKM for NM_008745, 19.30 +/− 4.45 FPKM for NM_001025074) (Fig. 1b).

**Figure 1.**
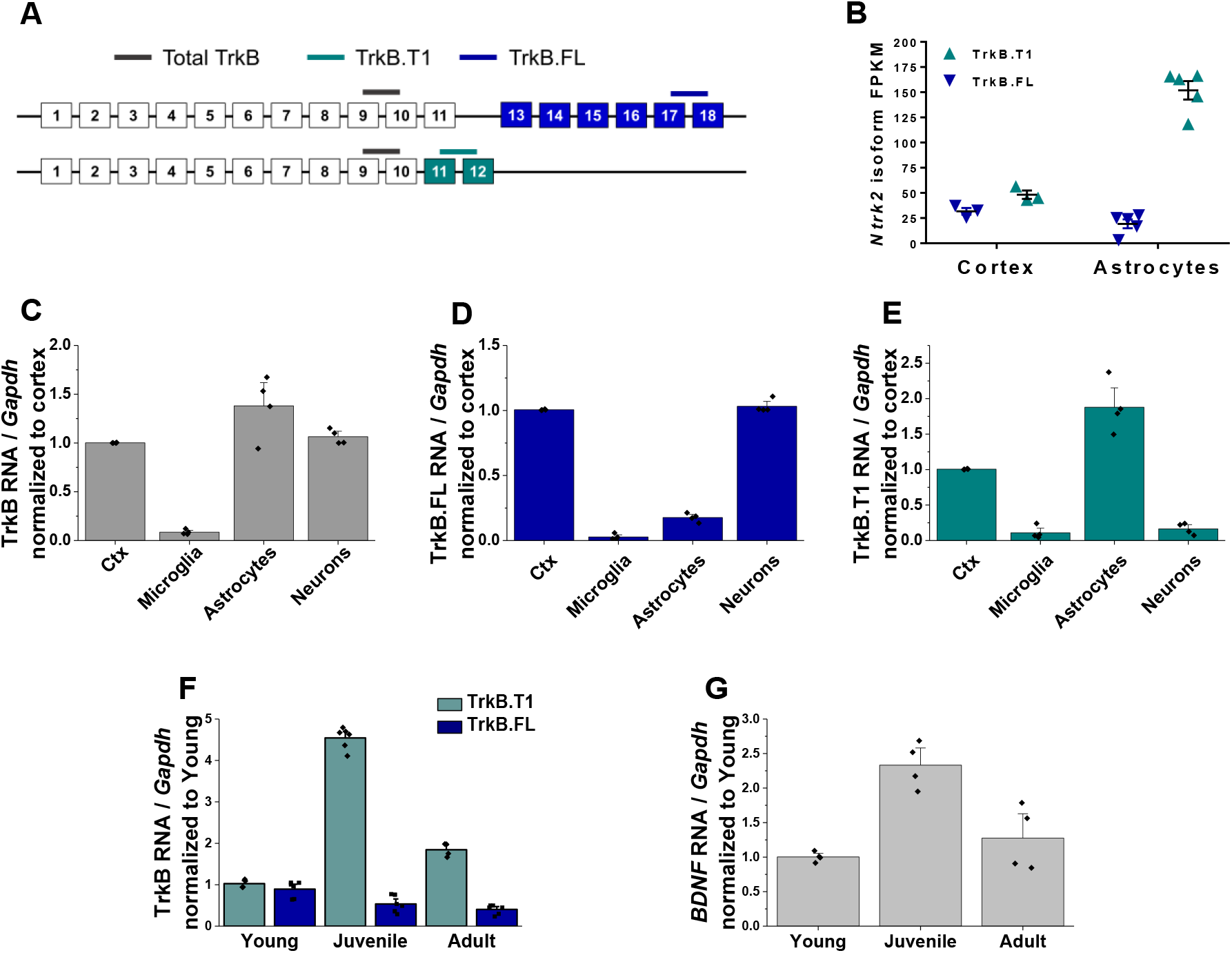
Astrocytes express high levels of truncated TrkB during astrocyte morphogenesis. **A** Cartoon representation of *Ntrk2* isoform exons and primers utilized. Total TrkB was probed with exons across both isoforms, with isoform-specific exons utilized to probe TrkB.Fl (blue) and TrkB.T1 (teal) **B** Total and isoform-specific *Ntrk2* RNA expression in astrocytes and whole cortex from juvenile animals. The majority of Ntrk2 isoform expression in astrocytes is attributed to the truncated TrkB.T1 receptor isoform. Quantitative PCR analysis of acutely isolated CNS populations from juvenile mice for **C** overall *Ntrk2*, **D** TrkB.FL, or E TrkB.T1 mRNA expression, normalized to matched whole cortex. **F** *Ntrk2* receptor isoform mRNA expression in astrocytes across development. Expression of TrkB.T1 is highest in juvenile animals, when **G** Bdnf mRNA is highest in cortical tissues. Data represented as mean +/− SEM, n = 3-6 animals.

This result prompted us to evaluate total, full length, and truncated *Ntrk2* mRNA expression in astrocytes relative to other CNS cell populations. Sequential isolation of oligodendrocytes, microglia, astrocytes, and neurons was performed as we have previously described (Holt and Olsen, 2016) in late juvenile mice. Cellular purity was confirmed via qPCR by evaluating cell type specific gene expression (Fig S1). Oligodendrocytes were excluded for subsequent analysis due to lack of cellular purity (Fig. S1). QPCR analysis of total and isoform-specific *Ntrk2* mRNA expression indicated total TrkB (primer detects both isoforms, grey in Fig 1A) was most highly expressed in astrocytes, relative to neurons or microglia. TrkB.FL is the predominant isoform expressed by neuronal populations (Fig. 1d). As indicated by the above RNA sequencing data, astrocytes predominately expressed the truncated TrkB.T1 expression (Fig. 1e). Expression of the truncated receptor is highest during the juvenile period (PND 28, Fig. 1f) relative to young (PND 8) and adult (PND 60) astrocytes when total availability of BDNF peaks in the cortex (Fig. 1g). Intriguingly, this time period correlates with the height of astrocyte morphological maturation.

### Novel serum-free primary astrocyte cultures

To test a direct effect of BDNF/TrkB on astrocyte morphogenesis we turned to a novel *in vitro* astrocyte culture system. Here, astrocytes were acutely isolated from postnatal day 3-6 pups utilizing a previously published MACS sorting technique (Holt and Olsen, 2016; Kahanovitch et al., 2018), with the modification of elution and plating in a serum-free, defined media (Fig. 2a). Cellular purity of the cultures was verified via qPCR, with mRNA levels of GFAP that are comparable to age-matched cortex but nearly undetectable levels of microglial, oligodendrocyte, OPC, and neuronal gene expression (Fig. 2b) at both 7 and 14DIV (Table 1). TrkB mRNA and protein expression was additionally verified, with similar levels of TrkB.T1 mRNA as age-matched *in vivo* astrocytes (Fig. 2c). Importantly, *in vitro* astrocytes exhibited a developmental upregulation of TrkB.T1 mRNA at 14DIV. Immunofluorescent co-staining of astrocytes with TrkB and GFAP revealed TrkB protein expression in astrocytes, with localization to astrocytic membrane (Fig. 1D).

**Figure 2.**
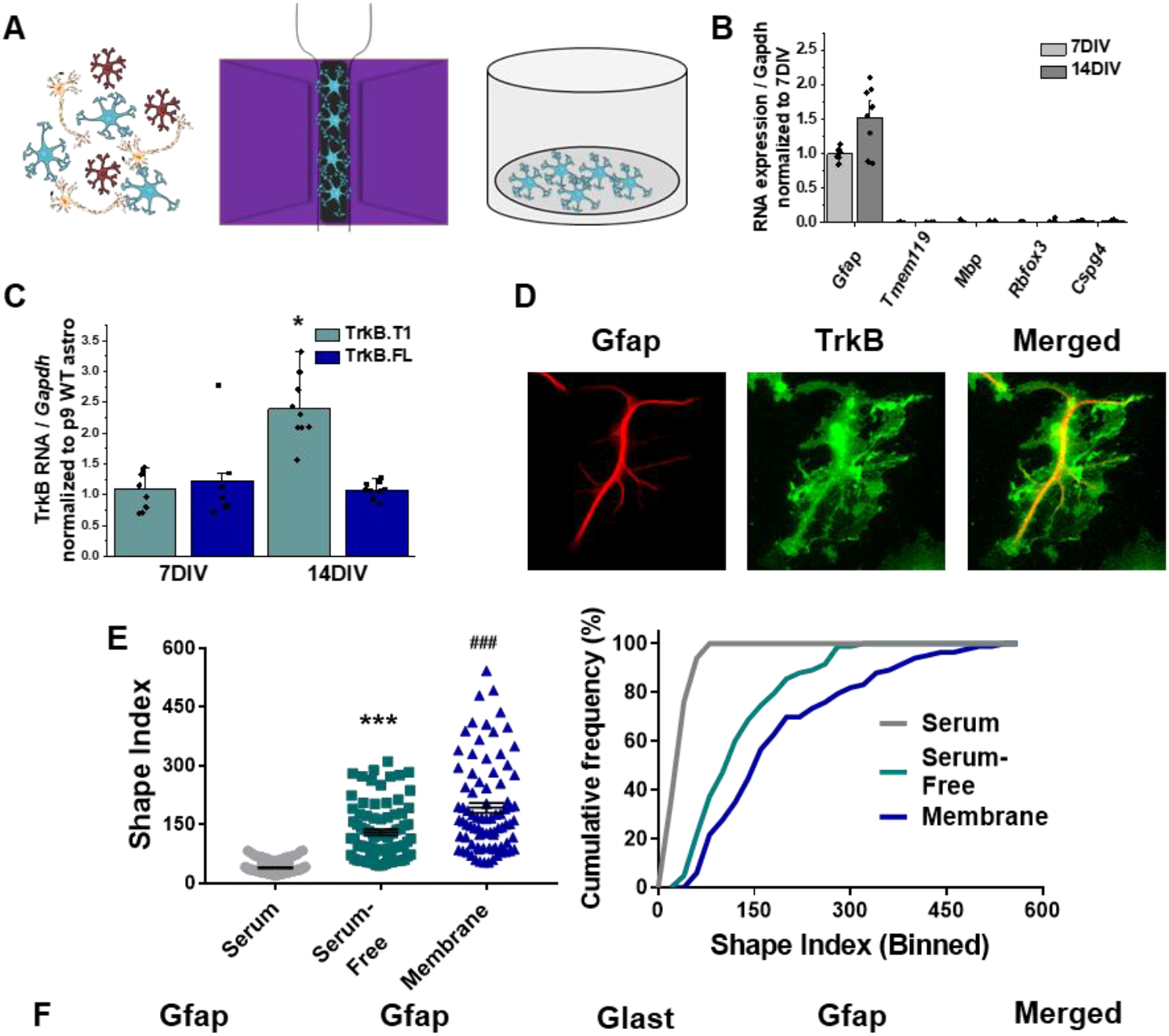
Novel serum-free primary astrocyte culture yields morphologically complex astrocytes. **A** Cartoon representation of magnetic separation of astrocytes for culture in serum-free, defined media. **B** Quantitative PCR data from cultured astrocytes at 7 and 14 DIV demonstrates purity of cultured cells. **C** mRNA expression of *Ntrk2* isoforms in cultured astrocytes compared to age-matched acutely isolated astrocytes demonstrates a developmental upregulation of TrkB.T1 expression. **D** Representative image of GFAP and TrkB immunofluorescence shows localization of TrkB to astrocytic membrane. **E** Shape index analysis and **F** cumulative frequencies of astrocytes cultured in serum-containing or serum-free media and membranous immunolabeling demonstrates increased cellular complexity in serum-free conditions. f Representative images of GFAP+ astrocytes cultured for 14DIV in the presence or absence of serum, and representatives images of membrane (Glast)/GFAP staining. Data represented as mean +/− SEM, n = 3-6 cultures, with 2 biological replicates per culture, *p < 0.05.

**Table 1.** Antibodies, AAV and pharmacological agents and catalog, catalogue, company and concentrations

| Reagent | Utilization | Company (Cat. No.) | Concentration / Time |
| --- | --- | --- | --- |
| Rb-GFAP | In vitro astrocyte morphology—branch processes | DAKO (Z0334) | 1:1000 / Overnight |
| Rb-Glast | In vitro astrocyte morphology—membrane | Abcam (ab416) | 1:500 / Overnight |
| Ms-Ezrin | In vitro astrocyte morphology—peripheral processes | Sigma (E8897) | 1:500 / Overnight |
| Gp-VGLUT1 | Excitatory presynapse | Millipore (AB5905) | 1:1000 / Overnight |
| Ms-PSD95 | Excitatory postsynapse | NeuroMab (75028) | 1:1500 / Overnight |
| Rb-TrkB (pan) | Visualization of full length and truncated TrkB protein | Millipore (07-225) | 1:1000 / 1 hour, WB<br>1:1000 / Overnight, IF |
| AAV2/5<br>GfaABC1D.mCherry | Astrocyte morphology – Sholl analysis | Addgene (#58909) | 2-3uL 3.67 x 10<sup>8</sup> |
| AAV2/5<br>GfaABC1D.lck-eGFP | Astrocyte morphology—Volume reconstruction | Addgene (#105598) | 2-3uL 2.3 x 10<sup>8</sup> |
| TrkB-Fc | BDNF scavenger | R&D Systems (# 688-TK-100) | 2ug |

Notably, culturing astrocytes in serum free media resulted in 3.43-fold more complex astrocyte as assessed by Shape Index of GFAP immunostained cells (t (198) = 13.39, p < 0.0001; Fig. 2e-f). A cumulative frequency analysis was performed to account for the large range in the datasets, and demonstrates a significant right shift in serum-free astrocytes (D = 0.75; p < 0.001). We additionally utilized a combination of Glast and Ezrin immunocytochemistry to demarcate both the membrane (Glast) and the fine, peripheral processes (Ezrin) in subsequent experiments. Glast— a membrane localized glutamate transporter—is highly expressed in astrocyte populations (Kondo et al., 1995) and Ezrin—a member of the ERM protein family—links the plasma membrane to the actin cytoskeleton, and has been previously demonstrated to be localized to peripheral astrocyte processes (Derouiche and Frotscher, 2001; Lavialle et al., 2011). The visualization and quantification of astrocyte membrane, which accounts for upwards of 85% of *in vivo* astrocytic volume, results in more physiologically relevant information. Unsurprisingly, comparison of Shape index quantification from membrane (Glast) staining revealed a 1.5 fold increased cellular complexity relative to intermediate filaments (GFAP) staining, (t(164) = 4.183, p < 0.001; Fig 2e-f).

### BDNF induces an increase in astrocyte morphological complexity

Given the high levels of astrocytic TrkB expression during a period of astrocyte morphological maturation we next evaluated a role for BNDF on astrocyte morphology. Astrocytes were isolated and cultured as described above, and experiments performed after 14 DIV. Wildtype (WT) astrocytes were exposed to 10ng, 30ng, or 100ng BDNF for 24 hours, followed by paraformaldehyde fixation. These concentrations were chosen based upon their previous use in investigating BDNF’s effects on neurons (Ji et al., 2005; Kline et al., 2010) and astrocytes (Ohira et al., 2007). Experiments confirmed that Glast and Ezrin targets did not change expression following BDNF exposure (SFig. 2a). Shape Index complexity analysis revealed BDNF-treated astrocytes showed a 2-fold increase in average astrocyte morphological complexity after exposure to 30ng BDNF (F(3, 214) = 7.047; p = 0.001; Fig 3a-c). Cumulative frequency analysis additionally demonstrated a right shift at 30ng BDNF (H(4) = 28.93; p = 0.006), indicating a BDNF-induced increase in astrocyte morphological complexity. We confirmed this finding with similar experiments performed in cells stained with GFAP to visualize astrocyte branch processes. SI quantification revealed a significant 1.4-fold increase in average astrocyte morphological complexity after exposure to 30ng BDNF (SFig. 2b-d). Given that 30ng BDNF exposure increased both astrocyte process and total cellular complexity, all following experiments were performed with this concentration.

**Figure 3.**
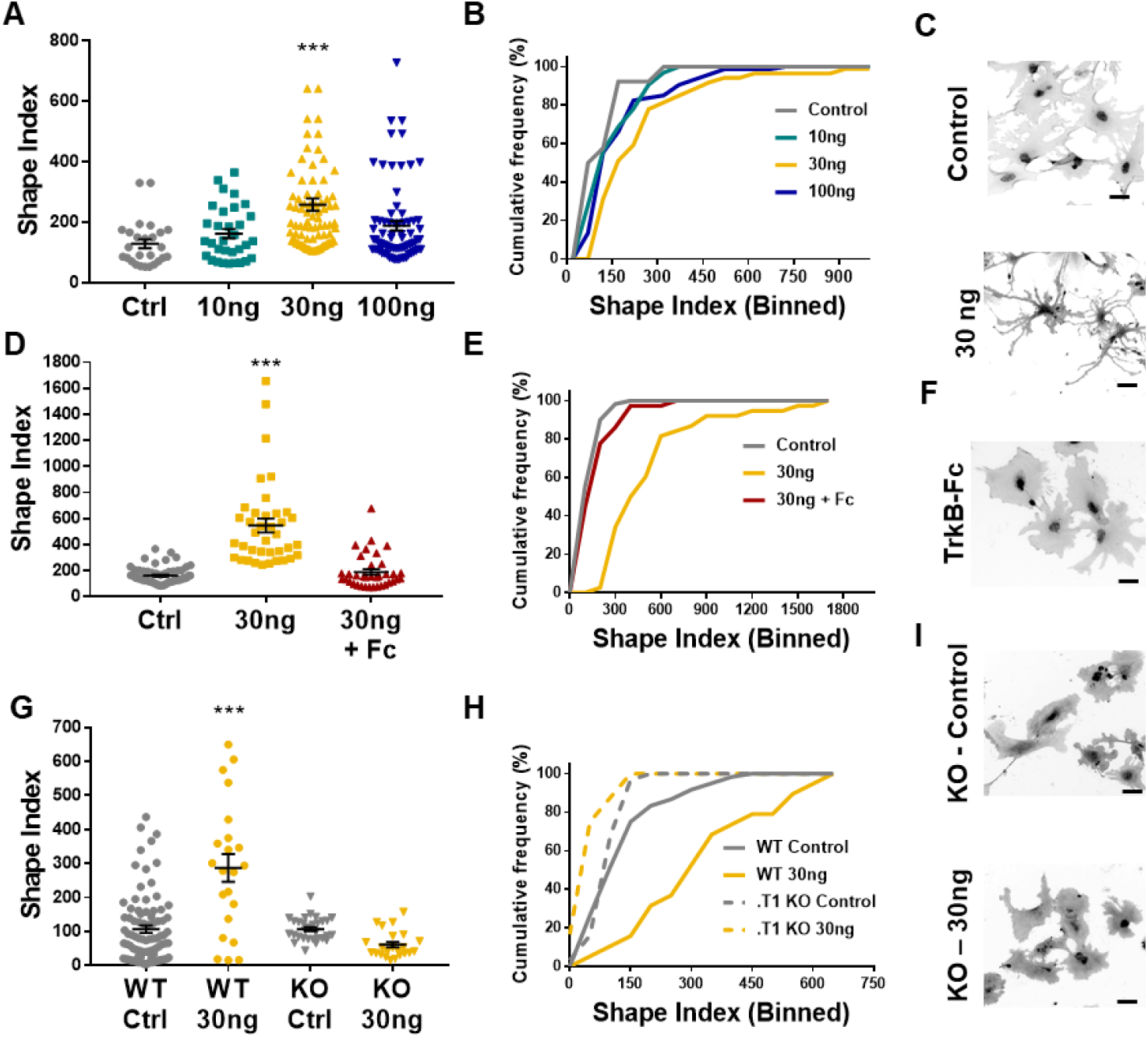
Astrocyte morphological complexity is sensitive to BDNF/TrkB.T1 signaling. **A** SI analysis and **B** cumulative frequencies following exposure to varying concentrations of BDNF for 24 hours. **C** Representative images of control and 30ng-treated astrocytes. **D, E** Complexity analysis of astrocyte morphology following scavenge of BDNF with TrkB-Fc, **F** representative image of TrkB-Fc scavenged astrocytes. **G-I** Morphological complexity analysis of cultured astrocytes lacking the TrkB.T1 receptor in the presence of BDNF. Data represented as mean +/− SEM, n = 3-6 cultures, with 2 biological replicates per culture. Scale bars indicate 20 microns. *p < 0.05, **p < 0.01, ***p < 0.0001.

### BDNFS effects are mediated via the truncated TrkB receptor

We next performed two loss of function experiments to ascertain the specificity of BDNF/TrkB.T1 effects on astrocytes. Current pharmacological TrkB receptor antagonists do not specifically target the truncated TrkB.T1 receptor. Therefore, we utilized TrkB-Fc to scavenge BDNF from the culture media. TrkB-Fc mimics the binding site of TrkB, allowing it to bind to BDNF and prevent BDNF binding to endogenously expressed TrkB receptors (Guo et al., 2012). WT 14DIV astrocytes were treated with 30ng BDNF, and one hour later 2ug TrkB-Fc additionally added. SI quantification of cellular complexity 24 hours later demonstrated that scavenging BDNF inhibited the increase in astrocyte morphological complexity (H = 70.41; p = .98; Fig. 3d-f).

To determine the necessity of TrkB.T1 receptor signaling in astrocytes we utilized a TrkB.T1 KO mouse model (Dorsey et al., 2006). We first validated specific loss of the truncated receptor in both tissue and astrocyte cultures. Both qPCR and western blot analysis of whole cortex and isolated astrocytes demonstrated loss of TrkB.T1, specifically (SFig. 3a, b). Importantly, western blot quantification demonstrated that TrkB.T1 KO mice do not exhibit a compensatory upregulation of the full length TrkB isoform (SFig. 3c). Note, in WT cortex, the band at the lower molecular weight, representing TrkB.T1 is expressed at higher levels than TrkB.FL, supporting our RNA sequencing data from whole cortex (58% vs. 42% of total TrkB, t(4) = 3.419; p = 0.03; Fig. 1B and SFig. 3C). Astrocytes were isolated and cultured from male pups as described above. Quantitative PCR and immunocytochemistry experiments demonstrated that these cultures do not express the truncated TrkB receptor at the mRNA or protein level (SFig. 3e-f). At 14DIV, WT and .T1 KO astrocytes were exposed to 30ng BDNF for 24 hours, and cellular complexity determined. As before, WT astrocyte SI indicated an increase in cellular complexity in response to BDNF (F(3, 155) = 21.66; p = .0001). However, SI quantification and cumulative frequency analysis revealed no difference in control- and 30ng-treated TrkB.T1 KO astrocyte cellular complexities (F(3, 155) = 21.66, p = 0.39; H (4, 136) = 40.88, p = 0.99, respectively; Fig. 3d-f). Our data, therefore, suggests that BDNF signaling through TrkB.T1 increases astrocyte morphological complexity at both the process and total membranous levels.

### In vivo loss of TrkB. T1 decreases astrocyte morphogenesis

The experiments above established that BDNF signaling through the TrkB.T1 receptor induces an increase in astrocyte morphological complexity in a simplified model system. We set out to examine astrocyte morphogenesis as an indicator of astrocyte developmental maturation in TrkB.T1 KO mice. Astrocyte morphology was examined in WT and TrkB.T1 knockout male animals at PND 14 and PND 28. Here the early time point represents a period in astrocyte development when astrocytes are considered morphologically immature (Bushong et al., 2004; Morel et al., 2014). Intraventricular injections of AAV2/5 GfaABC1D driven lck-GFP in postnatal day 0/1 pups allows for sporadic labeling of astrocytes throughout the brain (Fig 4a,b). Confocal z-stack images of layer II/III motor cortex astrocytes were acquired. Imaris surface reconstruction allows for the determination of the full astrocyte morphology, and is indicative of the amount of neuropil infiltration of the astrocyte peripheral processes (Morel et al., 2014). In line with previous reports, WT astrocyte volume increased by 1.75-fold between PND 14 and PND 28 (F(3,85) = 15.31, p < 0.001), indicative of normal morphological maturation (Fig. 4b) (Bushong et al., 2004; Morel et al., 2014; Stogsdill et al., 2017). In contrast, this increase in morphogenesis was lost in TrkB.T1 KO animals, with no significant difference between PND 14 and 28 astrocyte volumes (F(3, 85) = 15.31; p = 0.27; Fig. 4b). No difference between WT and TrkB.T1 KO astrocyte morphology was detected at PND 14 (F(3, 85) = 15.31; p = 0.86; n = 16 cells from 5 KO animals; n = 18 cells from 5 WT animals). By PND 28, TrkB.T1 KO astrocytes demonstrated a 30% reduction in volume compared to WT littermates (Fig 4b; F(3, 85) = 15.31; p = 0.003; n = 29 cells from 6 WT animals, n = 27 cells from 7 KO animals). Cumulative frequency analysis demonstrated a significant right shift between WT PND 14 and 28, indicating an increase in morphological complexity in wildtype astrocytes (H(4, 89) = 32.44; p < .001), with no difference between p14 and p28 TrkB.T1 KO astrocytes (H(4, 89) = 32.44; p = .728). We additionally performed a secondary analysis at p28, with an AAV driving cytosolic mCherry expression in astrocytes to better visualize process branching. Subsequent Sholl analysis revealed a decreased level of astrocytic branching (F(45, 495) = 1.55, p = 0.0149; SFig 4C) with no change in primary branch length (t(527) = −1.26, p = 0.21; SFig. 4D). Thus, our data suggests that BDNF signaling onto TrkB.T1 in astrocytes is an important pathway for normal astrocyte morphogenesis.

**Figure 4.**
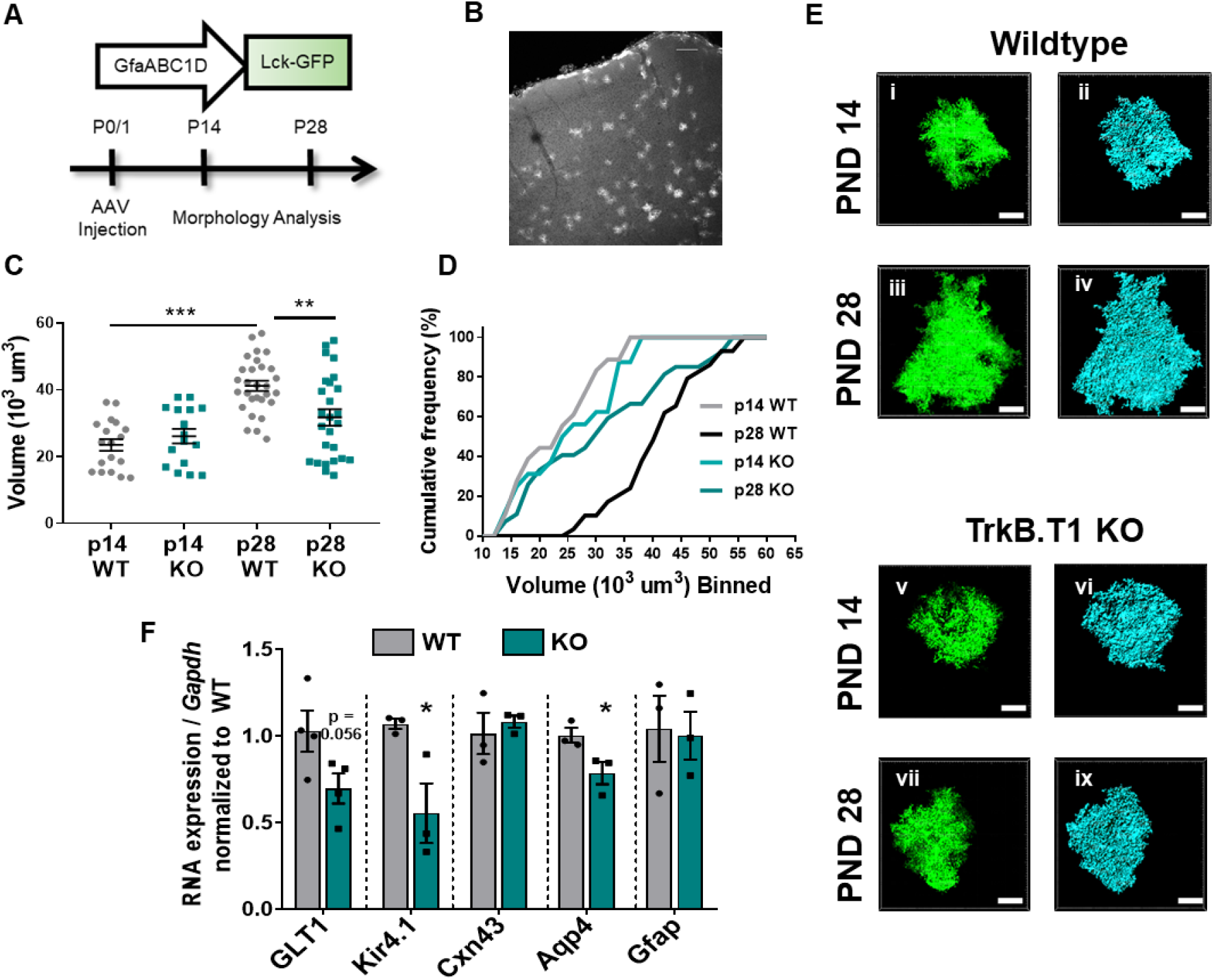
Loss of truncated TrkB.T1 *in vivo* leads to decreased astrocyte morphogenesis and an immature phenotype during maturation period. **A** Schematic of AAV injection and timeline of morphological analysis. **B** Representative image demonstrates sporadic, single astrocyte labeling in PND28 animals following AAV injections. Quantification of **C** astrocyte volume reconstruction and **D** cumulative frequency analysis demonstrates that prior to entering the morphological refinement phase (PND 14), wildtype and TrkB.T1 Ko astrocytes exhibit no difference in their morphological complexity. However, by the end of the maturation phase (PND 28), TrkB.T1 KO astrocytes fail to increase their volume. **E** Representative images of confocal (left, green) and Imaris surface reconstructions (right, blue) of astrocytes in (i-iv) wildtype and (v-ix) TrkB.T1 KO mice. **F** QPCR analysis of WT and .T1 KO PND25 astrocytes demonstrates decreased RNA expression of mature astrocytic genes Glt1, Kir4.1, and Aqp4, with no change in Cxn43 or Gfap expression. Data represented as mean +/− SEM, n = 5 - 7 animals for image analysis, with at least n = 3 cells per animal; n = 3 animals for qPCR analysis; *p < 0.05, **p < 0.01, ***p < 0.0001.

### *In vivo* loss of TrkB. T1 results in aberrant astrocytic gene expression

The period of astrocyte morphogenesis overlaps with differential gene expression in astrocytes in the developing cortex (Clarke et al., 2018; Molofsky and Deneen, 2015). We thus next examined genes located perisynaptically that are associated with mature astrocyte functions (Clarke et al., 2018; Molofsky and Deneen, 2015; Nwaobi et al., 2014; Regan et al., 2007). TrkB.T1 KO and WT littermate astrocytes were acutely isolated in juvenile males (PND 25) as described above. QPCR analysis revealed decreased RNA expression in specific gene sets differentially regulated in astrocytes (Clarke et al., 2018): Kir4.1 (51.6%, t(4) = 2.943; p = 0.04) and Aqp4 (21.7%, t(4) = 2.807; p = 0.04), with trending decrease in Glt1 (33.1%, t(6) = 2.242; p = 0.056) relative to WT littermate controls (Fig. 4F). No significant difference was observed in genes that display no change in cortical development: Cxn43 (t(4) = 0.5491, p = 0.612) and Gfap (t(4) = 0.171, p = 0.87) (Fig. 4F). Thus, *in vivo* loss of TrkB.T1 results in dysregulated expression of genes associated with mature astrocyte function.

Astrocyte morphological maturation also overlaps with neuronal synaptogenesis and refinement (Bushong et al., 2004; Freeman, 2010; Morel et al., 2014), and astrocytic contributions to synaptogenesis represent an intense investigative area of astrocyte biology (Allen and Eroglu, 2017) Therefore, we next probed the above TrkB.T1 KO astrocytes for known astrocyte synaptogenic factors SPARCL1/hevin and SPARC (ref) via qPCR. We observed a 28.55% decrease in SPARCL1/hevin RNA expression in .T1 KO astrocytes compared to WT littermates (t(5) = 4.061; p = 0.0097). Interestingly, we additionally observed a 56.16% increase in hevin-antagonist SPARC (t(6) = 4.061; p = 0.0215) in .T1 KO astrocytes compared to WT littermates. These experiments together suggest a dysregulation of astrocyte gene expression in TrkB.T1 KO animals, with consequences on mature astrocyte functions.

### Potential role for BDNF/TrkB.T1 astrocyte signaling on neuronal synapse development

The above experiment suggests dysregulated astrocyte synaptogenic factor expression in .T1 KO astrocytes. In order to query if BDNF signaling onto astrocytes contributes to neuronal synapse development, we turned to a novel neuron-astrocyte co-culture model system. This system allows us to specifically and directly evaluate the interplay between maturing astrocytes and neurons. To this end, wildtype cortical neurons were cultured from p0/1 pups, and WT or TrkB.T1 KO astrocytes subsequently layered on top after 3 days of recovery. After 8DIV, cells were paraformaldehyde fixed and confocal images of excitatory (VGlut1+/PSD95+) synapses acquired. Excitatory synapses were evaluated due to extensive literature detailing astrocytic contributions to excitatory synaptogenesis (reviewed in (Allen and Eroglu, 2017). Puncta Analysis quantification of colocalization of pre- and post-synapses (Ippolito and Eroglu, 2010; Stogsdill et al., 2017) revealed that, as expected, excitatory synapse numbers were increased in the presence of WT astrocytes (F(2, 37) = 5, *p* = 0.013) which did not occur in the presence of TrkB.T1 KO astrocytes (F(2, 39) = 5, p = .846). Notably, further analysis revealed this effect appears to be influenced by differential consequences on pre- or postsynaptic sites. We observed a similar increase in the number of presynaptic puncta in excitatory synapses compared to WT astrocytes (F(2, 42) = 5.44, p = 0.799) with a significant reduction in post-synaptic partners (F(2, 42) = 5.44, p = 0.015). This led to a significant reduction in co-localized and presumably functional synapses (Fig. 5a, b) (Ippolito and Eroglu, 2010; Stogsdill et al., 2017). Of note, these findings correspond with the decreased hevin / increased SPARC expression observed in TrkB.T1 KO astrocytes. These studies suggest, at least in a simple model system, TrkB.T1 astrocytes do not support normal excitatory synaptic development.

**Figure 5.**
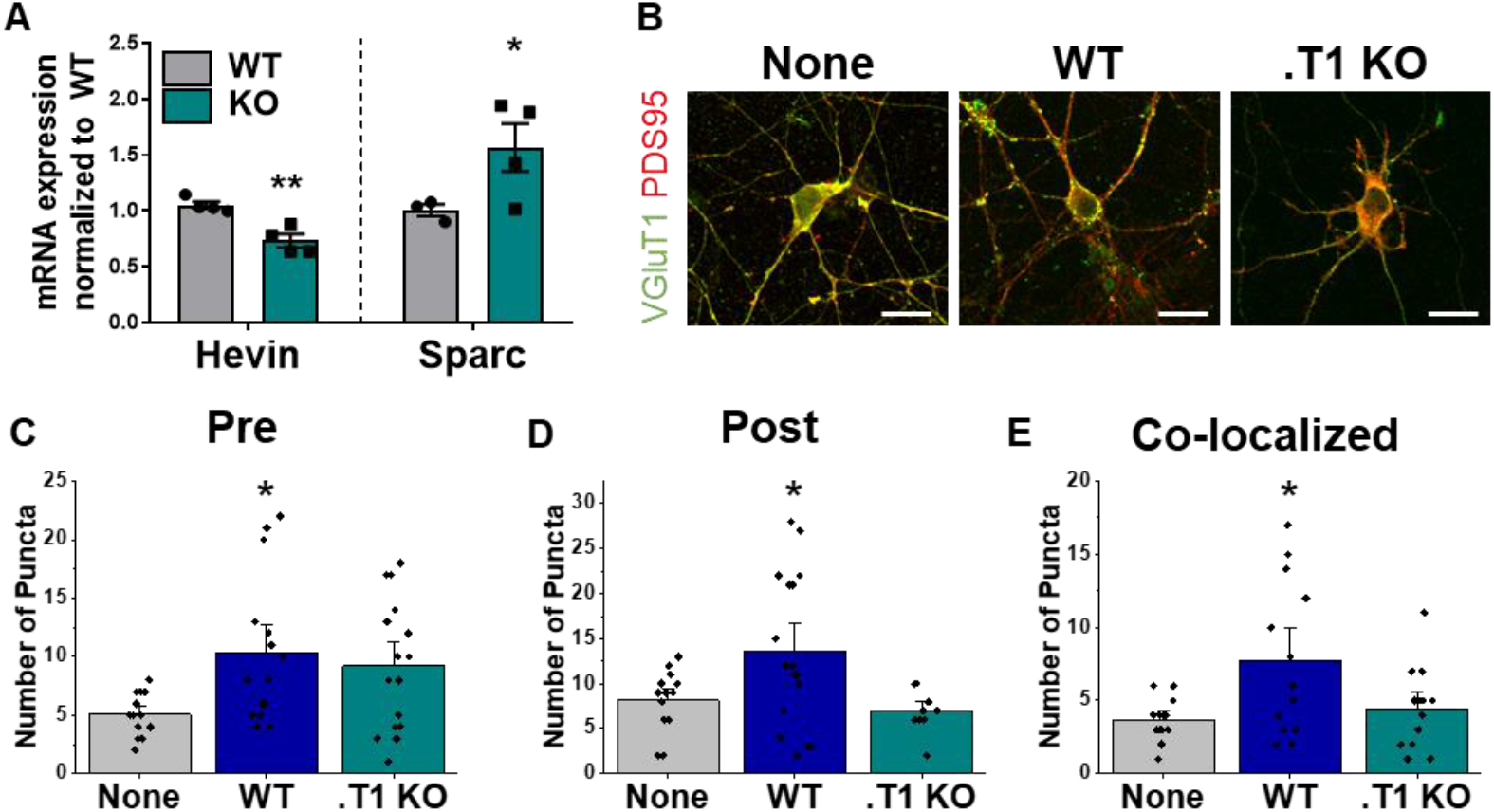
Astrocytes lacking TrkB.T1 receptor are unable to support neuronal synaptogenesis. **A** QPCR analysis of acutely isolated PND25 WT or TrkB.T1 KO astrocytes revealed aberrant RNA expression of known astrocyte synaptogenic factors Hevin and Sparc in .T1 KO astrocytes compared to wildtype. **B** Representative images of neurons cultured in the presence of none, wildtype, or TrkB.T1 KO astrocytes. Excitatory synapses were visualized with VGlut1+/PSD95+ colocalization, with inhibitory synapses visualized with VGat+/Gephyrin+ colocalization. Quantification of B pre-synaptic, **C** post-synaptic, and **D** colocalized, functional synapses of cultured neurons. Data represented as averages +/− SEM, n = 34 animals for qPCR; N = 4 cultures, with at least n = 5 cells per culture. *p < 0.05, **p < 0.01; * = compared to no astrocyte control; # = compared to WT astrocytes.

### Discussion

Herein we have demonstrated for the first time that BDNF is involved in the maturation of a non-neuronal cell type. Within cortical astrocytes, TrkB.T1 receptor expression is in the top 20 of all protein-coding transcripts, with the highest expression during astrocyte morphological refinement. We developed and utilized a novel astrocyte culture paradigm to demonstrate that BDNF induces an increase in astrocyte morphological complexity, which is dependent upon the TrkB.T1 receptor. Importantly, *in vivo* astrocyte morphology is less complex with loss of the TrkB.T1 receptor. In particular, the lack of TrkB.T1 prevented normal morphogenesis during the time of astrocyte morphological refinement, between p14 and p28, indicating BDNF/TrkB.T1 signaling may drive astrocyte morphological maturation. Furthermore, BDNF signaling onto astrocytes appears to have consequences on neuronal synapse development, with decreased numbers of both excitatory and inhibitory synapses in the presence of TrkB.T1 KO astrocytes. These findings have broad implications, given the wealth of neurological disorders within which BDNF and, increasingly more often, astrocytes are implicated.

BDNF’s role in CNS growth and maturation has been intensely studied, with a particular emphasis on the full length receptor and neuronal function. Here, we demonstrate that astrocytes express the highest levels of *Ntrk2* over other CNS cellular populations. Publicly available RNA sequencing databases corroborate our data (Zhang et al., 2014), and, in fact, additionally demonstrate that human astrocytes express the highest levels of *Ntrk2* (Kelley et al., 2018; Zhang et al., 2016). In concordance with previously published results (Rose et al., 2003), we found that neurons predominately express the full-length isoform of TrkB. While *in vitro* and *in situ* hybridization studies have demonstrated expression of the truncated TrkB isoform in glial populations (Park and Poo, 2013; Rose et al., 2003), we demonstrate for the first time that *in vivo* astrocytes express developmentally regulated levels of TrkB.T1, with the highest expression occurring during juvenile stages. This upregulation of TrkB.T1 is mediated by an intrinsic, cell-autonomous mechanism, as cultured astrocytes demonstrated similar upregulation between 7 and 14DIV.

Truncated TrkB.T1 lacks the canonical tyrosine kinase signaling domain, and therefore has historically been presumed to act in a dominant negative capacity to prevent overactivation of BDNF signaling pathways (Fenner, 2012; Klein et al., 1989; Middlemas et al., 1994). We here have shown that BDNF induces an increase in astrocyte morphological complexity, and this effect is lost in TrkB.T1 KO astrocytes. Therefore, our data suggests an intrinsic and direct mechanism of action. Supporting this, *in vivo* dysfunction of astrocytic TrkB.T1 has been implicated in mediating neuropathic pain and motor dysfunction following spinal cord injuries (Matyas et al., 2017). BDNF application to cultured astrocytes resulted in a PLCy-IP3R mediated rise in calcium with kinetics different from cultured neurons (Rose et al., 2003). Additionally, the TrkB.T1 receptor has been found to co-immunoprecipitate with a RhoGTPase inhibitor, RhoGDIa in primary astrocyte cultures, and inhibition of RhoA increased the area occupied by cultured astrocytes (Ohira et al., 2005). Given RhoGTPases’s known roles in regulating astrocytic cytoskeletal dynamics (Zeug et al., 2018), this presents a potential mechanism by which BDNF increases astrocyte morphological complexity in astrocytes. It is unclear how BDNF is processed by astrocytes following binding to TrkB.T1. However, our data suggests TrkB.T1 plays an active, direct role in astrocyte biology.

Little is known regarding the mechanisms governing astrocyte morphogenesis and maturation. Thus far, three mechanisms have been identified. In drosophila, FGF signaling through the Heartless receptor determines astrocyte domain size and infiltration into the neuropil (Stork et al., 2014). Pharmacological and genetic manipulations of mGluR5 in astrocytes reduced the developmental increase in astrocytic volume between PND14 and 21 (Morel et al., 2014). Similarly, loss of neuroligins in astrocytes and/or their neuronal neurexin partner reduced astrocyte morphogenesis in the visual cortex by PND21. Based on our findings, we propose BDNF signaling through truncated TrkB.T1 receptor as a novel mediator of astrocyte morphological maturation. This is supported by our evaluation of known developmentally regulated and mature astrocyte functional markers, Kir4.1 and Glt1. Similar to others (Morel et al., 2014; Stogsdill et al., 2017), we found that wildtype astrocytes exhibited a 1.75-fold increase in volume during astrocyte morphological maturation. However, TrkB.T1 KO astrocytes did not exhibit normal morphogenesis. No difference in astrocyte volume between young and mature ages in TrkB.T1 KO animals was found, indicative of a failure to properly undergo morphogenesis and maturation, and resulted in a 30% decrease in astrocyte volume compared to mature wildtype littermates. An important limitation to our data is the utilization of a global TrkB.T1 knockout mouse model to assess *in vivo* astrocyte morphology. However, given that in astrocyte cultures, whereby the loss of the receptor is cell-type specific, TrkB.T1 KO astrocytes do not morphologically respond to BDNF. Additionally, qPCR analysis of microglia, astrocytes, and neurons at the time of astrocyte morphological maturation indicates that TrkB.T1 receptor expression is largely confined to astrocyte populations. We therefore make the argument that within the CNS, the majority of effects within the global TrkB.T1 KO mouse may be attributable to astrocytes. Regardless of the in vivo cell-type specificity, our data cumulatively highlights BDNF signaling through astrocytic TrkB.T1 receptor as a mediator of astrocyte morphological maturation.

Peripheral astrocyte processes, which are dramatically increased in complexity during the morphological refinement phase, facilitate many known astrocyte biological functions including enwrapment of synapses. Astrocytes actively contribute to synaptogenesis through release of astrocyte derived factors such as hevin (Kucukdereli et al., 2011), thrombospondins (Christopherson et al., 2005), SPARC (Kucukdereli et al., 2011), and glypicans 4/6 (Allen et al., 2012). Astrocyte enwrapment of synapses is additionally known to be regulated by neuronal activity, and can stabilize synapses following LTP (Bernardinelli et al., 2014). We found decreased expression of hevin, and increased expression of its antagonist SPARC. As TrkB.T1 KO astrocytes demonstrate decreased morphological complexity and dysregulated astrocyte synaptogenic factor expression, we additionally investigated how BDNF signaling onto astrocytes may impact neuronal synapse number. To this end, we developed a novel astrocyte-neuron co-culture paradigm. Utilization of MAC sorting technique allows for separation and subsequent combination of different cellular subtypes, genotypes, and ages within cultures. We posit that this technique will be useful to many for investigations of cell-to-cell communication. We found neurons cultured in the presence of .T1 KO astrocytes exhibited decreased numbers of overall excitatory post-synaptic elements and an overall reduction in numbers of co-localized pre and post-synaptic elements. These findings corroborate the aberrant RNA expression observed, as hevin and SPACR control the structural formation of synapses, particularly at the postsynaptic side (Jones et al., 2011; Kucukdereli et al., 2011). Notably, thus far, no reports have explored interactions between BDNF and known astrocyte-mediated synaptogenic factors. It is also noteworthy that experiments revealing astrocyte-mediated synaptogenesis are often performed in neuron-astrocyte co-cultures with BDNF added as a supplement (Allen et al., 2012; Johnson et al., 2007; Pfrieger and Barres, 1997; Ullian et al., 2001). Future studies are needed to elucidate BDNF’s role in astrocyte-mediated synaptogenesis and refinement.

We demonstrated that scavenging BDNF from the media within an hour of exposure prevented the increase in cellular complexity 24 hours later, suggesting that BDNF must be actively present to elicit an increase in astrocyte morphological complexity. This experiment is particularly interesting given that following synaptogenesis, BDNF secretion from neurons is largely targeted to synaptic zones and is secreted in an activity-dependent manner (Park and Poo, 2013). While outside of the scope of this paper, the influence of BDNF on astrocyte morphological complexity may indeed extend into activity-dependent maintenance of astrocyte morphology and enwrapment of synapses. One study suggests that this may indeed occur, as siRNA knockdown of TrkB.T1 in adult rats leads to decreased ability of cortical astrocytes to modulate their morphology in response to neuronal activity (Ohira et al., 2007). These results also highlight that BDNF may be necessary for the maintenance of astrocyte morphology in adulthood.

Here we demonstrate BDNF’s receptor, TrkB.T1, is highly enriched in cortical astrocytes, particularly during the period of astrocyte morphological maturation, and that BDNF/TrkB.T1 signaling in astrocytes plays a critical role in astrocyte morphogenesis and may play a role in proper astrocyte maturation. Furthermore, proper neuronal synaptogenesis was lost with deletion of the TrkB.T1 receptor in astrocytes. Our studies suggest that BDNF/TrkB.T1 signaling is a novel unexplored pathway in the role of astrocytes in synapse development. Given the role of aberrant synapse development in neurological dysfunction, our results herein suggest astrocyte BDNF/TrkB.T1 signaling may contribute to neurodevelopmental disorders in which BDNF signaling is implicated.

## Methods

### Animals

All experiments were performed according to NIH guidelines and with approval from the Animal Care and Use Committee of the University of Alabama at Birmingham and Virginia Polytechnic Institute and State University. All animals were maintained on a 12 hour light/dark cycle (lights on at 9pm, lights off at 9am) with food and water available *ad libitum*. Every effort was made to minimize pain and discomfort. Wild-type and *TrkB.T1*^−/−^ and wild-type littermate mice (Dorsey et al., 2006) C57/B6 mice were used for these experiments. *TrkB.T1*^−/−^ mice were a generous gift from Dr. Lino Tessarollo.

### Cortical Dissection and Dissociation

Briefly, mice (young, postnatal day 7 +/− 1 days (PND 7), late juvenile mice (PND 28 +/− 3 days (PND 28) or adult mice (PND 60+/− 10 days (PND 60) were anesthetized via CO_2_ and decapitated. Whole cortex was microdissected in ice-cold ACSF (120mM NaCl, 3.0 mM KCl, 2mM MgCl, 0.2mM CaCl, 26.2mM NaHCO3, 11.1 mM glucose, 5.0mM HEPES, 3mM AP5, 3mM CNQX) bubbled with 95% oxygen. Tissue was minced into 1mm^3^ pieces and dissociated for 15-30 minutes using Worthington Papain Dissociation Kit. Tissue was subsequently triturated until a single-cell suspension was achieved and filtered through a 70μM filter.

### Astrocyte Isolations

Astrocytes were acutely isolated as previously described (Holt and Olsen, 2016; Kahanovitch et al., 2018; Stoica et al., 2017). Following dissociation, microglia and myelin were first removed from the cell suspension. Cells were incubated for 15 minutes at 4°C with 15μL of Miltenyi Biotec’s Myelin Removal Kit and Cd11b^+^ MicroBeads. The suspension was then applied to a prepped LS column, washed three times, and the flow-through collected. This flow through was subsequently used to isolate astrocytes utilizing Miltenyi Biotec’s ACSA-2^+^ MicroBead kit. The cell suspension (in 150uL 0.5% fatty-acid free BSA in PBS) was incubated at 4°C for 15 minutes with 15-20μL FcR blocker, followed by a 15 minute incubation with 15-20μL ACSA-2 microbeads. Cells were applied to a prepped LS column. Astrocytes were eluted from the LS column after three washes, with 5mL buffer and the supplied plunger.

### Sequential CNS population isolations

Cells were acutely isolated as previously described (Holt and Olsen, 2016). Following dissociation, oligodendrocytes were isolated first with a 10 minute incubation with 15uL Myelin+ microbeads. Cells were applied to a prepped LS column, and washed 3x. All flow through was collected and utilized to isolate the subsequent cellular populations. Oligodendrocytes were eluted from the LS column after three washes, with 5mL buffer and the supplied plunger. Microglia were isolated next, with a 10 minute incubation with 15 uL Cd11b+ microbeads. Cells were applied to a prepped LS column, and washed 3x. As before, all flow through was collected and utilized to isolate the next cellular population. Microglia were eluted from the LS column after three washes, with 5mL buffer and the supplied plunger. Astrocytes were subsequently isolated as described above. Finally, neuronal populations were isolated using Neuronal isolation kit. The flow through was again collected and used to isolate neurons utilizing Miltenyi Biotec’s Neuron Isolation Kit. The cell suspension was incubated with 20μL biotinylated antibodies for 10 minutes at 4°C, followed by a 15 minute incubation with 20μL anti-biotin microbeads. Cells were applied to a prepped LD column. Neurons were collected in the flow through of two washes.

### RNA isolation and qPCR

Total RNA was isolated using Ambion’s PureLink RNA Mini Isolation kit according to the manufacturer’s instructions. RNA samples designated for RNA Sequencing were eluted in 30 μL filtered, autoclaved Mill-Q water. Subsequently, 2ng of RNA was reverse transcribed into cDNA using BioRad’s iScript kit or BioRad’s iScript SuperMix. All cDNA was normalized to 350 or 500ng (for BDNF mRNA assays) following conversion. The relative mRNA expression levels were determined using real-time quantitative PCR by General Taqman PCR master mix and TaqMan specific probes (Table 1). Relative mRNA expression levels were determined by the ddCt method, with each normalization indicated where appropriate.

### RNA Sequencing

RNA samples were tested for quality on the Agilent Tapestation 2200 (Agilent Technologies, Santa Clara, CA). The NEB Next rRNA Depletion Kit (NEB #E6310X) was used to process 250 ng of total RNA. RNA-Seq libraries (400 bp) were created using the NEBNext Ultra II Directional RNA Library Prep Kit for Illumina (NEB #E7760L). Samples were individually indexed using the NEBNext Multiplex Oligos for Illumina (NEB #E6609S). Adapter ligated DNA was amplified in 13 cycles of PCR enrichment. Libraries were quantified with the Quanti-iT dsDNA HS Kit (Invitrogen) and qPCR. Library validation was performed on the Agilent 2200 Tapestation. Independently indexed stranded cDNA libraries were pooled and sequenced for 150 cycles with the Illumina NovaSeq 6000 S2 Kit. All samples were sequenced at 85-90 million read depth, paired-end 2 x 150 bp, and in reverse-stranded orientation.

### Bioinformatics analyses

Initial analyses (raw reads processing through read alignment) were run in the University of Alabama at Birmingham’s Cheaha High Performance Computing (HPC) cluster environment. Raw RNA-Seq reads were concatenated (per R1 and R2 fastq read, respectively) and quality trimmed using Trim Galore! Version 0.4.3. Sequence quality of trimmed reads was inspected using FastQC (version 0.11.15). The STAR aligner (version 2.5.2) (Dobin et al., 2013) was used in the basic two-pass mode to align the trimmed reads to the iGenomes UCSC mm10 mouse genome. BAM files were sorted by coordinate, and indexed using SAMtools (Li et al., 2009) (version 1.3.1). To examine general gene expression levels, a gene counts table was created using featureCounts (Liao et al., 2014) (release 1.5.2) and used as input for DESeq2 (Love et al., 2014) (version 1.16.1) in the RStudio environment (version 3.4.1). Genes with a row sum less than 10 were excluded prior to differential gene expression analysis. Normalized counts were extracted for each biological replicate to calculate the average normalized counts per respective gene. For transcript expression analysis, the STAR-aligned BAM files were processed in the University of Alabama at Birmingham Galaxy platform (Afgan et al., 2016) using Stringtie (Pertea et al., 2016) (Galaxy tool version 1.3.3.1) as described in the recommended workflow, with minor modifications: 1) the reverse strand option was selected and 2) the iGenomes UCSC mm10 genome was used as the reference guide assembly data set for the first Stringtie run. *Ntrk2* transcript expression levels (FPKM) were extracted from the second Stringtie run’s Assembled Transcripts output files per respective biological replicate. All detailed scripts used for these analyses are available upon request.

### Protein extraction and immunoblotting

Proteins were extracted by homogenizing samples in lysis buffer (1% sodium dodecyl solfate (SDS), 100mM Tris(hydroxymethyl)aminomethane (Tris) buffer, pH 7.5), supplemented with protease and phosphatase inhibitors (Sigma), followed by two rounds of sonication for seven seconds. Lysates were subsequently centrifuged for 5 minutes at 16,000 x*g*. ThermoScientific’s Peirce BCA assay was utilized to determine protein concentrations. Proteins were heated to 60°C for 15 minutes with 2x loading buffer (100mM Tris, pH 6.8, 4% SDS, in Laemmli-sodium dodecyl sulfate, 600mM B-mercaptoethanol, 200mM Dithiothreitol (DTT), and 20% glycerol). Equal amounts of protein per sample (5 or 10ug) was loaded into a 4-20% gradient precast mini-PROTEAN TGX gel (Bio-Rad) and proteins were separated with 200V in 1x running buffer (24.76 mM Tris base, 190mM glycine, 0.1% SDS). Proteins were transferred to a nitrocellulose membrane using the Trans-blot turbo system (Bio-Rad), mixed molecular weight protocol (2.5A, 25V for 7 min), followed by 1 hour blocking with LI-COR blocking buffer at a 1:1 ratio with TBS. Primary antibodies, including concentration and incubation times, are given in Table 2. All secondary antibodies were LI-COR, and incubated at 1:10,000 for 1 hour at room temperature. Imaging was performed on a LI-COR Odyssey machine on both the 680 and 800 channels.

**Table 2.**
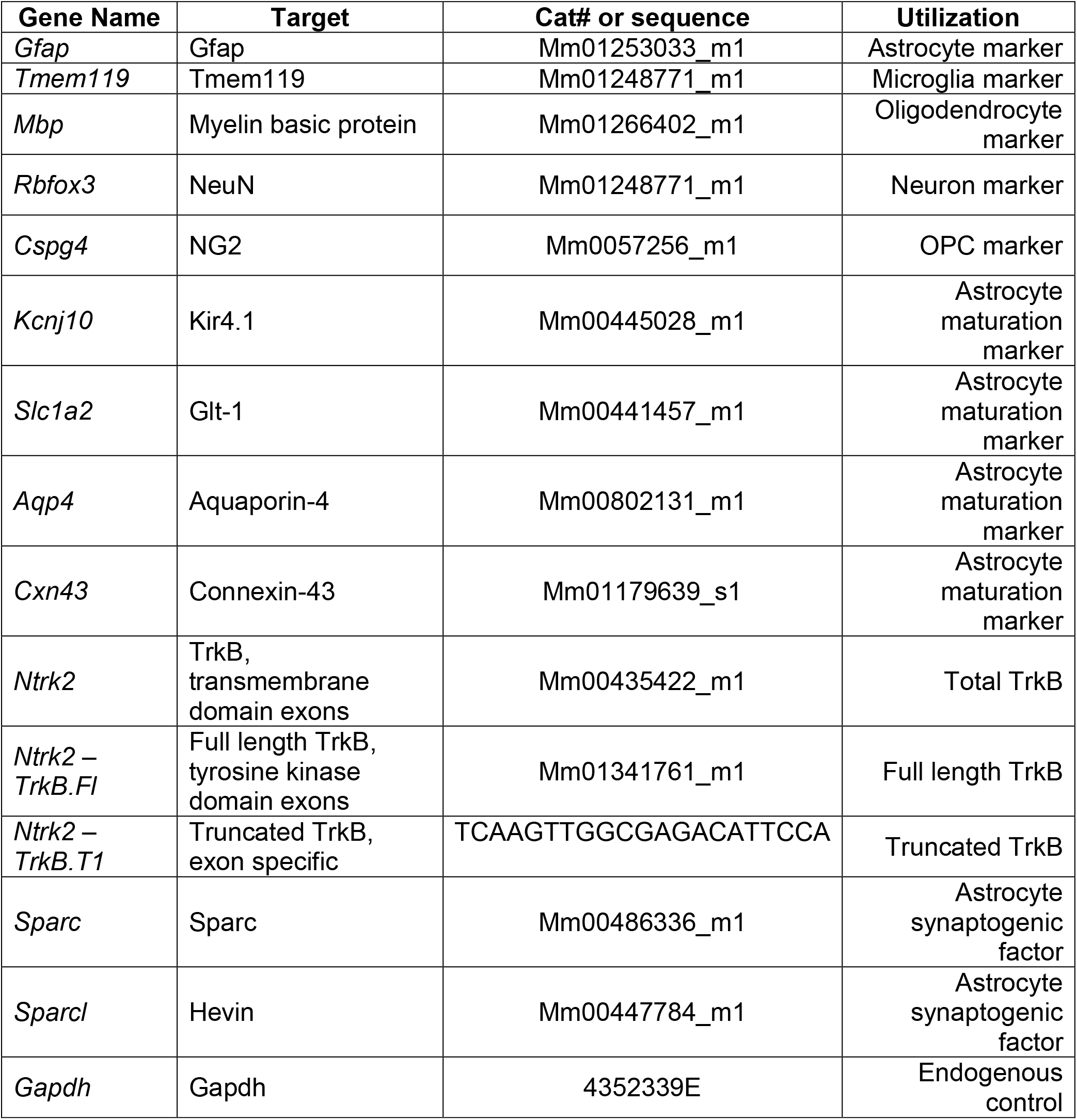
QPCR primers and catalog numbers utilized.

### Serum-free primary astrocyte culture

Astrocytes were isolated from postnatal day 3-6 pups as described above and previously (Kahanovitch et al., 2018). Following elution, astrocyte cell number was determined, and 0.75-1.0 x 10^5^ cells were plated on 13mm glass coverslips in a 24 well plate. The coverslips were poly-l-ornithine treated and laminin-coated. Astrocytes were maintained in serum-free, defined media (50% Neurobasal media, 50% MEM, 1mM sodium pyruvate, 2mM glutamine, and 1x B27). On the first day post-plating, fresh media was added. On the third day post-plating, a complete media change was performed. Subsequent media changes occurred every 3-4 days. Astrocytes were collected at 7 and 14 days in vitro (DIV).

### Primary astrocyte culture experiments

Primary astrocyte cultures were utilized at 14 DIV for experiments. Exogenous BDNF (Promega) was applied in warmed media to a final concentration of 0ng, 10ng, 30ng, or 100ng for 24 hours. For scavenger experiments, a final concentration of 2ug of TrkB-Fc (R&D Systems) was added to the wells in warmed media 60 minutes post-BDNF exposure. Equal volumes of warmed media as the TrkB-Fc condition was additionally added to controls.

### Primary astrocyte culture immunofluorescence

Astrocytes were fixed at 15 DIV, after experiments described above. First, pre-warmed paraformaldehyde (PFA) was added to the culture dish to a final concentration of 2% PFA, and incubated for 5 minutes at 37°C. This initial step was utilized to preserve any fine peripheral astrocyte processes that might be sensitive to cold temperatures. After incubation, cells were washed with cold PBS, followed by fixation with 4% PFA for 15 minutes at room temperature. Subsequently, cells were incubated for 1 hour in blocking buffer (10% goat serum, 0.3% Triton-X in PBS). Astrocyte filaments were visualized with GFAP, total membrane with Glast and Ezrin. Following primary antibody incubations, AlexaFlour 488, 546, and 647 were utilized to visualize the primary antibodies with 1 hour incubations. Prior to image acquisition, the experimenter was blinded to experimental conditions. Fluorescent images were acquired with an Olympus VS-120 system or Nikon A1 confocal.

### Primary astrocyte culture morphology analysis

The complexity of astrocytes following experiments described above was determined by utilizing the Shape Index, given as perimeter^2^ / area −4π (Holt and Olsen, 2016; Matsutani and Yamamoto, 1997). A perfect circle results in an index of 1, and increasingly complex cells have correspondingly larger indexes. Area and perimeter of the cells were determined manually using ImageJ 1.52b version software. Prior to quantification, experimenter was blinded to experimental conditions.

### In vivo astrocyte morphological analysis

Astrocytes were fluorescently labeled via AAV2/5-driven mCherry or lck-GFP. Lck-GFP virus— pAAV.GfaABC1D.PI.Lck-GFP.SV40—was a gift from Baljit Khakh (Addgene viral prep # 105598-AAV5). AAV2/5 GfaABC1D.mCherry was obtained from Vector Biolabs. Postnatal day 0-1 pups were intraventricularly injected with 2-3μL 2.3 x 10^8^ virus following hypothermia-induced anesthesia. The injection site was determined following (Chakrabarty et al., 2013; Kim et al., 2014; Shen et al., 2001), with equidistance between the bregma and lambda sutures, 1mm lateral from the midline, and 3mm depth. Hamilton 10μL syringes and 32G needles were used. Animals were collected at PND14 and PND28-30 (referred to PND28 in manuscript). At time of collection, animals were deeply anesthetized with peritoneal injections of 100mg/kg ketamine and intracardially perfused with PBS, followed by 4% PFA for 20 minutes. Brains were post-fixed for 72 hours, and subsequently sliced on Pelco Easislicer microtome at 100μM sections. Experimenter was blinded to animal genotypes prior to image acquisition and analysis. Layer II/III motor cortex astrocytes were imaged on a Nikon A1 confocal with 40x oil immersion lens (OFN25) and 3x digital zoom. Z-stacks were acquired with 0.225 μM step sizes. Laser power and gain were adjusted for each individual astrocyte. Z-stacks were 3D reconstructed on Imaris x64 9.0.2, and surface reconstruction utilized to estimate astrocyte volume. Sholl analysis was additionally performed in Imaris. Following surface reconstruction of astrocyte branch processes, the Filaments function was utilized followed by subsequent Sholl analysis function. Prior to quantification, experimenter was blinded to experimental conditions.

### Primary neuron culture

Neurons were cultured from p0-1 mouse pups according to Beaudoin et al. 2012 and as described above with modifications. In brief, following cortical dissociation, microglia and oligodendrocytes were first removed with 10μL incubation with Cd11b+ and Mbp+ microbeads for 10 minutes. The flow through was collected and used to further isolate neuronal populations utilizing Miltenyi Biotec’s Neuron Isolation Kit. The cell suspension was incubated with 10μL biotinylated antibodies for 10 minutes at 4°C, followed by a 10 minute incubation with 15μL anti-biotin microbeads. Cells were applied to a prepped LS column. Neurons were collected in the flow through of two washes. Neuronal cell number was determined, and 0.75-1.0 x 10^5^ cells were plated on 13mm glass coverslips in a 24 well plate. The coverslips were poly-l-lysine treated and laminin-coated. Neurons were maintained in neuronal maintenance media (Beaudoin et al., 2012) (Neurobasal media, 2mM l-glutamine, and 1x B27). On the first day post-plating, 2uM of araC was added to reduce non-neuronal contamination. On the second day post-plating, a media change was performed to remove araC. Subsequent media changes occurred every 3-4 days. At 3DIV 0.75-1.0 x 10^5^ WT or T rkB.T1 KO astrocytes from p5 pups were plated on top of neurons. Cells were collected at 8-9 DIV for synapse quantification.

### Neuron synapse quantification

Cells were fixed at 8-9 DIV, after experiments described above. Fixation was performed as described above. Subsequently, cells were incubated for 1 hour in blocking buffer (10% goat serum, 0.3% Triton-X in PBS). Excitatory synapses were visualized with presynaptic marker VGLUT1 and with postsynaptic marker PSD95. Following primary antibody incubations, AlexaFlour 488 and 647 were utilized to visualize the primary antibodies with 1 hour incubations at 1:500. These secondaries were chosen for their excitation/emission spectrum, which demonstrate no overlap. Therefore, co-localization analysis of pre- and post-synaptic markers can be utilized with confidence of true co-localization. Prior to image acquisition, the experimenter was blinded to experimental conditions. Confocal images were acquired on Nikon A1 confocal with 40x objective and 3 digital zoom. Care was taken to ensure each individual neuron imaged was equidistance from other neurons and astrocytes. Co-localization, and therefore synapse number, was determined utilizing Puncta Analysis FIJI plug-in (Ippolito and Eroglu, 2010; Stogsdill et al., 2017).

### Statistical analysis

To determine statistical significance, Origin and Graphpad Prism were utilized. All data is represented as mean +/− SEM, with n’s indicated where appropriate. D’Agostino-Pearson normality test was performed to determine the normality distribution of each data set, and outliers were determined via GraphPad Prism’s ROUT method. Student’s t-tests were performed, or Mann Whitney U tests for nonparametric data, for all data in which only one comparison was needed. One-way ANOVAs, or Kruskal-Wallis test for nonparametric data, followed by Tukey’s post-hoc test performed for all data with multiple comparisons.

## Acknowledgements

The authors wish to thank Lara Ianov and CIRC Neurodevelopmental Bioinformatics Iniative for their help in processing the RNA-sequencing and Lina Tessorallo and colleagues for the kind gift of the TrkB.T1 KO mouse model. This work was funded by the National Institutes of Health NINDS R01NS075062 (M.O.) and F31NS100259 (L.M.H.).

