## Supplemental Figures for "BDNF/TrkB.T1 signaling is a novel mechanism for astrocyte morphological maturation"

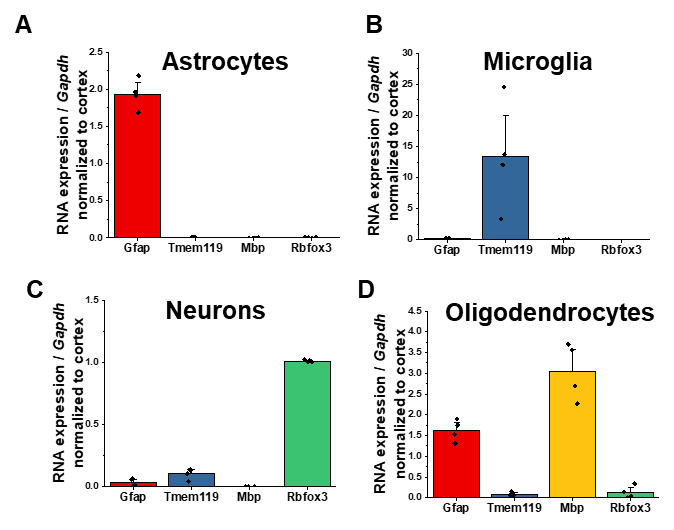

**Supplemental Figure 1**. Validation of isolated CNS populations. Quantitative PCR data of **A** astrocytes **B** microglia, **C** oligodendrocytes, and **D** neurons isolated from juvenile, PND25 animals. For microglia, astrocytes, and neurons, enrichment of cell-type specific genes with depletion of other cellular genes demonstrates isolation of pure cellular populations. Significant enrichment of Gfap was found in oligodendrocytes. Data represented as mean +/- SEM, n = 4 animals.

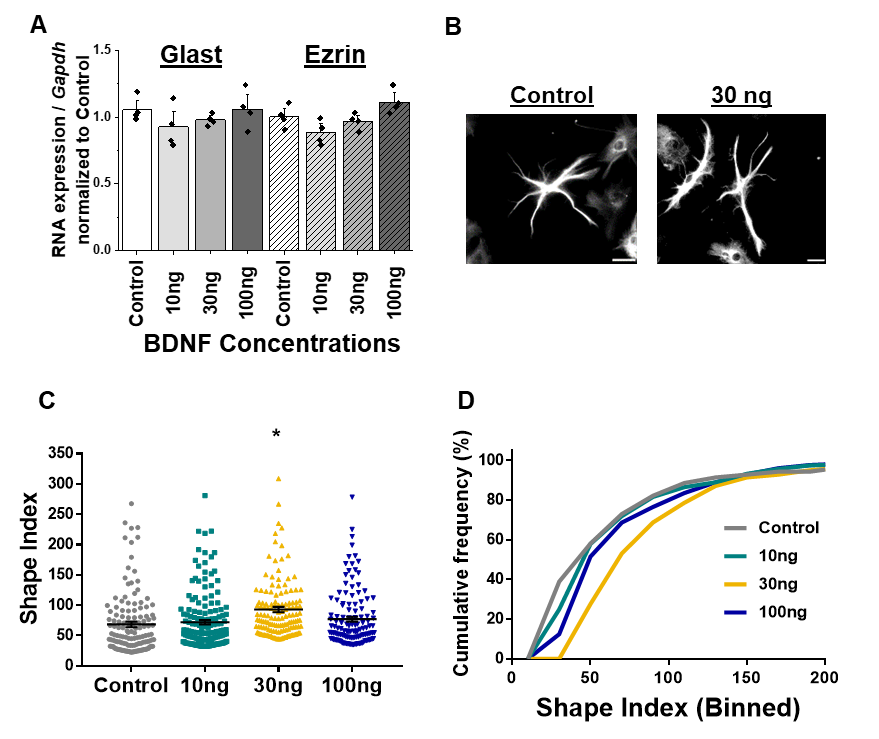

**Supplemental Figure 2**. Wildtype astrocytes increase process branch complexity in response to BDNF. **A** Quantification PCR validation that Glast and Ezrin expression does not change in response to varying concentrations of BDNF for 24 hours. **B** Representative images of GFAP+ processes in control- and 30ng-treated astrocytes. **C** Shape index quantification and **D** relative frequency curves demonstrate an increase in astrocyte process branch complexity in response to 30ng BDNF. Data represented as mean +/- SEM, n = 3-6 cultures, with 2 biological replicates per culture, and 3-5 random ROIs per image. Scale bars indicate 20 microns. *p < 0.05.

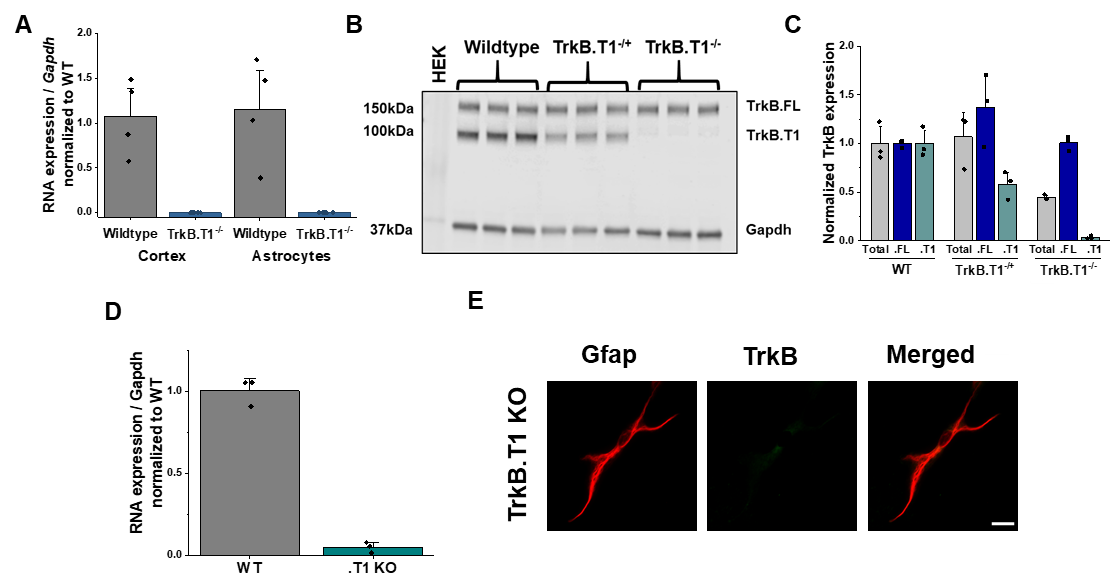
**Supplemental Figure 3**. Validation of truncated TrkB isoform specific deletion in TrkB.T1 KO mice. **A** Quantitative PCR data of whole cortex and isolated astrocytes from wildtype and TrkB.T1 KO animals demonstrates depletion of TrkB.T1 mRNA compared to wildtype levels. **B** Representative western blot and **C** quantification demonstrates no compensatory upregulation of the TrkB.FL isoform. **D** Quantitative PCR data of TrkB.T1 KO cultured astrocytes at 14 DIV demonstrates loss of TrkB.T1 mRNA expression. **E** Representative images of Gfap and TrkB immunofluorescence in 14 DIV TrkB.T1 KO astrocytes. Data represented as averages +/- SEM, n = 3-6 cultures, with 2 biological replicates per culture.

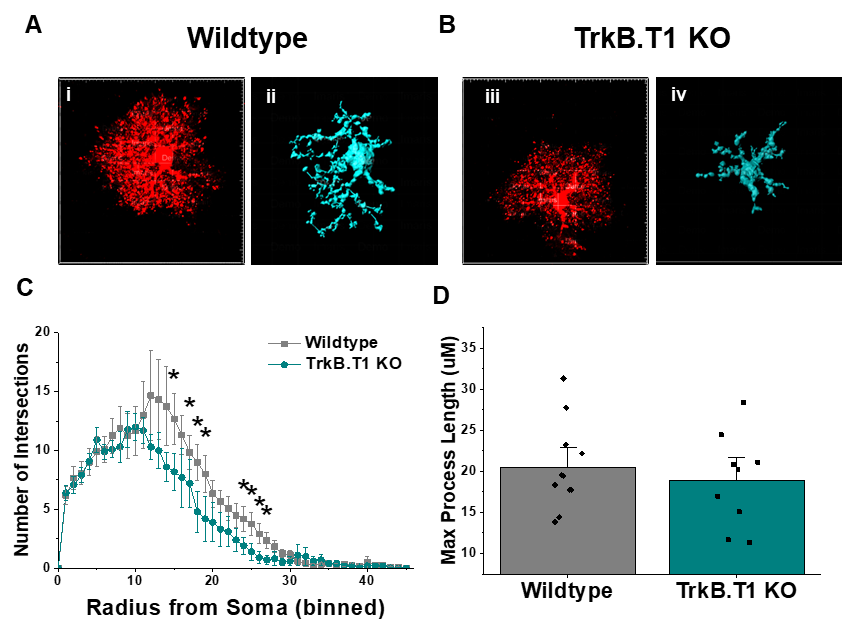

**Supplemental Figure 4**. Sholl analysis quantification of PND29 astrocytes. **A, B** Representative confocal images (**i**, **iii**) and Imaris reconstruction of primary branches (**ii**, **iv**) in wildtype and TrkB.T1 KO astrocytes. **C** No difference in max process length of astrocyte primary branches was found, with **D** decreased number of branch intersections in TrkB.T1 KO compared to wildtype littermates. Data represented as mean +/- SEM, n = 3 animals; *p < 0.05, **p < 0.01, ***p < 0.0001.
